# Encapsulated cell technology delivers ciliary neurotrophic factor to promote JAK/STAT-dependent photoreceptor survival in retinal degeneration

**DOI:** 10.64898/2026.07.27.740168

**Authors:** Yasuaki Iwama, Leia Laughlin, Sarah Harkins-Perry, Sarah Giles, Ava Maeyama, Kerstin Traxler, Mieke van Daelen, Roberto Bonelli, Kohji Nishida, Martin Friedlander, Marin L. Gantner, Kevin T. Eade

## Abstract

Sustained trophic factor delivery via Encapsulated Cell Technology (ECT) is a powerful new class of therapeutics with broad potential for targeted treatment. Intravitreal delivery of ciliary neurotrophic factor (CNTF) via the ECT, NT-501, is a first-in-class therapy that slows the progression of macular telangiectasia type 2 (MacTel). Despite its clinical efficacy, key questions remain regarding its mechanism of action, including whether other implant-derived factors contribute to therapeutic benefit and how optimal dosing should be determined. Resolving these issues is critical for optimizing NT-501 in MacTel and guiding the development of ECT-based therapies for other diseases. We evaluated the biological activity of implant-derived cytokines on retinal tissue, using long-term NT-501 intravitreal implants in rabbits alongside human retinal organoid (hRO) models treated with NT-501–conditioned medium (NT-501–CM). Then, using a MacTel-specific photoreceptor degeneration model in hROs, we showed NT-501–CM significantly reduced photoreceptor cell death, and this protective effect was abolished by either CNTF-neutralizing antibodies or JAK inhibitor. We also established a therapeutic dose-response relationship linking NT-501–derived CNTF levels to JAK/STAT3 activation and photoreceptor protection. These findings directly connect ECT-derived CNTF exposure with JAK/STAT3-mediated photoreceptor protection in human retinal tissue and suggest an optimal concentration range for efficacy.

## Introduction

Encapsulated cell technology (ECT) enables sustained, controlled delivery of therapeutic proteins *in vivo* and has emerged as a promising therapeutic strategy for chronic diseases such as neurodegenerative disorders (1–3). Among candidate neuroprotective factors, ciliary neurotrophic factor (CNTF) has attracted considerable interest because of its neuronal survival–promoting activity across the nervous system (4–6). In chronic neurodegenerative conditions, sustained neuroprotective signaling may be required to counter progressive neuronal loss (7, 8). The retina is a highly specialized extension of the central nervous system in which neuronal degeneration causes irreversible visual impairment, making it an important target for CNTF-mediated neuroprotection. Given the short half-life of CNTF (9), maintaining therapeutic activity remains a major challenge, making ECT a suitable strategy for sustained CNTF delivery in the retina.

Preclinical studies in animal models have demonstrated that exogenous CNTF delivery can protect retinal neurons and preserve retinal function, including attenuation of photoreceptor cell (PRC) degeneration (10–14). These protective effects have been associated with activation of intracellular signaling pathways (JAK/STAT, MAPK/ERK, and PI3K/Akt), regulating transcriptional responses for neuronal survival, apoptosis, and proliferation (8, 15). However, mechanistic insights have largely derived from acute and supraphysiological CNTF paradigms in simplified experimental systems that do not fully recapitulate sustained CNTF exposure or the extracellular environment associated with device-mediated delivery.

ECT–based NT-501 was developed as a first-in-class platform for sustained intravitreal CNTF delivery, using implanted cells engineered to continuously secrete CNTF into the vitreous cavity (16, 17). Building on the therapeutic potential of CNTF in retinal degenerative diseases, NT-501 has been evaluated across multiple chronic retinal and optic neuropathies (7, 18–22). Notably, in macular telangiectasia type 2 (MacTel), a progressive macular degenerative disease with limited treatment options (23, 24), NT-501 has been shown to significantly slow disease progression in clinical trials (22), and was recently approved for clinical use in MacTel. Importantly, CNTF delivered via NT-501 is secreted from engineered cell lines which likely release additional secreted factors, creating a complex signaling environment distinct from recombinant CNTF exposure alone. How this clinically relevant exposure paradigm influences intracellular signaling and PRC survival in the human retina remains unclear.

To address this gap, we developed an integrated experimental framework to define how NT-501 regulates intracellular signaling and PRC survival. We combined long-term NT-501 implantation in rabbit eyes with short-term NT-501–conditioned medium (NT-501–CM) exposure in human induced pluripotent stem cell (iPSC)-derived retinal organoids (hROs) to benchmark this human tissue system against *in vivo* vitreoretinal responses. Using transcriptomic and targeted proteomic analyses with TUNEL-based cell death assays, we examined whether NT-501 exposure activates JAK/STAT-associated survival signaling and how these responses relate to PRC survival. Finally, we tested whether these signaling responses are required for PRC protection in an hRO model of MacTel-specific PRC degeneration. This approach defines a mechanistic link between clinically relevant CNTF exposure and JAK/STAT–associated survival signaling in human retinal tissue, and establishes a framework for identifying therapeutic exposure ranges relevant to NT-501 efficacy.

## Results

### 1. Long-term NT-501 implantation exerts minimal metabolic effects while activating JAK/STAT signaling in the rabbit retina

To characterize tissue responses to long-term implantation of NT-501 *in vivo*, we implanted the device into one eye of each rabbit, using the contralateral eyes as controls. The devices remained implanted for 6 months, corresponding to the duration of the first follow-up visit in clinical trials (22), and the retinas were then harvested for bulk RNA sequencing (RNA-seq) and metabolomic analysis.

We utilized RNA-seq on rabbit retinas treated with NT-501 or control to broadly characterize the effects of long-term NT-501 exposure (5 biological replicates). RNA-seq quantified 10,347 genes, of which 10,282 were mapped to human orthologs for downstream pathway analyses. Differential expression analysis identified 1,084 upregulated and 833 downregulated genes (adjusted *P* < *0.05*; 1,917 genes in total; Figure 1A). Hallmark enrichment analysis revealed significant enrichment of immune-related pathways, including IFN-γ and IFN-α response, allograft rejection, and complement, as well as the IL-6–JAK/STAT3 signaling pathway (Figure 1B and Supplemental Figure 1A). KEGG analysis showed a similar overall trend, including enrichment of JAK/STAT signaling (Supplemental Figure 1B). Consistent with this, multiple genes associated with JAK/STAT signaling (red dots in Figure 1A) were significantly upregulated in the volcano plot. We also observed that genes in the KEGG phototransduction pathway were significantly depleted (Supplemental Figure 1B). The upregulation of JAK-STAT and downregulation of phototransduction genes, both known targets of CNTF (5), are highly indicative of an active CNTF response in the rabbit retinas treated with NT-501.

**Figure 1.**
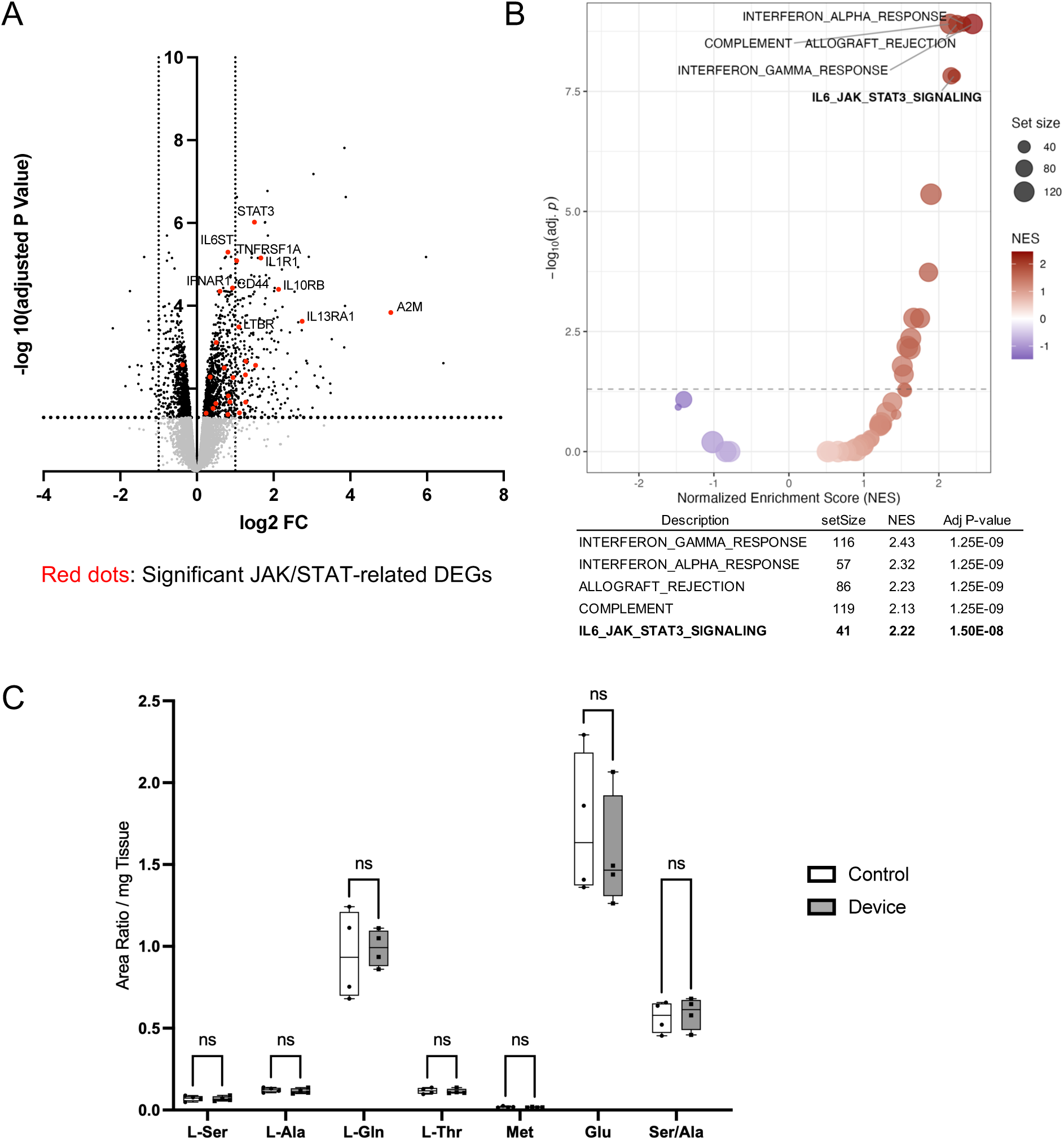
Effects of the long-term NT-501 implantation on retinal tissue in rabbits. (A) Volcano plot of differentially expressed genes in rabbit retinas following NT-501 implantation compared with contralateral control eyes, with genes mapped to human orthologs. Black dots indicate differentially expressed genes (FDR < 0.05), and red dots highlight JAK/STAT-related genes identified in enrichment analysis. (B) Gene set enrichment analysis of Hallmark gene sets showing enrichment in NT-501–treated rabbit retinas relative to contralateral control eyes. Each point represents a gene set, plotted by normalized enrichment score (NES) and –log10(adjusted P value). Top 5 significantly enriched gene sets ranked by adjusted P value are shown. (C) Quantification of metabolomic analysis showing mean area ratios normalized to tissue mass for L-serine (L-Ser), L-alanine (L-Ala), L-glutamine (L-Gln), L-threonine (L-Thr), methionine (Met), glutamate (Glu), and serine-to-alanine ratio (Ser/Ala). Statistical significance was determined using multiple paired t tests with Holm–Šídák correction for multiple comparisons. ns, not significant.

Because reduction of serine abundance leading to dyslipidemia is one of the possible disease mechanism of MacTel (25, 26), we next performed targeted metabolomic analysis of rabbit retinas to determine if the effects of NT-501 involve alterations in retinal metabolic profiles, either through direct release of serine or through cytokine signaling. Targeted metabolomic analysis demonstrated that retinal serine-related amino acid profiles, including serine-to-alanine ratios, were not significantly changed in implanted eyes compared with contralateral control eyes (Figure 1C). These findings suggest that long-term NT-501 implantation does not substantially alter the abundance of metabolites related to MacTel.

### 2. hROs treated with NT-501**–**CM have similar transcriptional response as rabbit retinas with long-term NT-501 implant

To determine the effects of NT-501 on human retinal tissue and establish a model for manipulation of NT-501–derived cytokines, we next treated hROs with NT-501–CM. To generate conditioned media, NT-501 was cultured under human endothelial serum-free medium, used as the control medium (Ctrl), and the supernatant, defined as NT-501–CM, was collected (See Methods; CM-1 through CM-5) containing 21.6 ± 4.29 ng/mL of CNTF (range, 17.7-27.2 ng/mL; see Methods). To investigate the NT-501–derived complex cytokine environment, multiplex cytokine analysis was performed on CM-1 through CM-5. Compared with Ctrl, all NT-501–CM samples showed markedly elevated levels of IL-8, IL-13, PDGF-AA, VEGF-A, CXCL16, SDF-1α+β, and MCP-1; IL-6 was likewise elevated in CM-1 through CM-4 but remained undetectable in CM-5 (representative CM-1 profile shown in Figure 2A). No other markers differed consistently between Ctrl and the NT-501–CM samples (Supplemental Table 1).

**Figure 2.**
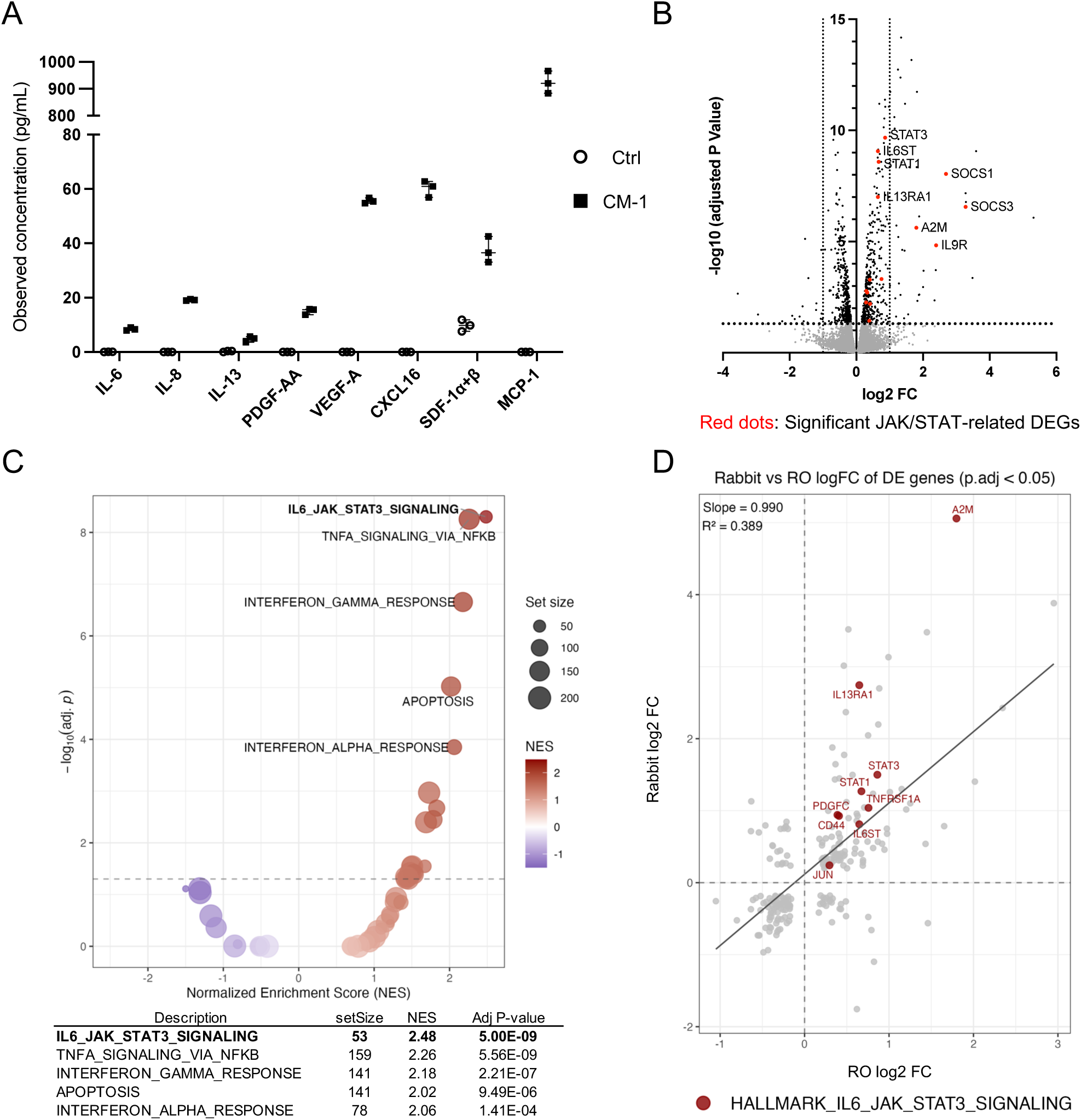
Composition and biological effects of NT-501–conditioned medium (NT-501–CM) on hROs. (A) Multiplex cytokine profiling of CM-1. Representative CM-1 profile showing selected cytokines that were consistently elevated in NT-501–CM samples compared with Ctrl. (B) Volcano plot of differentially expressed genes (DEGs) in hROs treated with NT-501–CM (CM-1) for 24 hours compared with Ctrl medium. Black dots indicate DEGs (FDR < 0.05), and red dots highlight JAK/STAT-related genes identified in enrichment analysis. (C) Gene set enrichment analysis of Hallmark gene sets showing enrichment in NT-501–CM relative to Ctrl after 24 hours of treatment. Each point represents a gene set, plotted by normalized enrichment score (NES) and –log10(adjusted P value). Top 5 significantly enriched gene sets ranked by adjusted P value are shown. (D) Comparison of differential gene expression responses between NT-501–implanted rabbit retina and CM-1-treated hROs. Each point represents a significantly DEG (adjusted P value < 0.05), plotted by logFC in rabbit retina (y-axis) versus logFC in hROs after 24 hours of NT-501–CM treatment (x-axis). Red dots indicate genes belonging to the HALLMARK_IL6_JAK_STAT3_SIGNALING gene set. The solid line represents linear regression, with the corresponding slope and R² values shown.

We next treated hROs with NT-501–CM where NT-501–CM was diluted to set the CNTF concentration at physiologically relevant levels, 100 pg/μL, approximately double the concentration observed in patient vitreous in clinical trials (19). As a negative control, hROs were cultured in Ctrl medium. Transcriptomic analysis by bulk RNA-seq was performed on hROs from each group after 24 hours of treatment (8 hROs per condition, 3 biological replicates). Differential expression analysis identified 462 upregulated and 395 downregulated genes (adjusted *P < 0.05*; 857 genes in total; Figure 2B). Hallmark gene set enrichment analysis (GSEA) revealed significant enrichment of IL-6–JAK/STAT3 signaling, along with IFN-γ and IFN-α responses and TNF-α signaling via NF-κB, in NT-501–CM-treated samples compared with Ctrl-treated samples (Figure 2C and Supplemental Figure 2A). KEGG analysis showed a similar overall trend, including enrichment of JAK/STAT signaling, while phototransduction-related pathways were negatively enriched (Supplemental Figure 2B).

To compare the *in vivo* effects of long-term NT-501 implantation in rabbit retinas with those observed *in vitro* using hROs, we performed a cross-species comparison of differential gene expression profiles. Among the 10,276 rabbit genes mapped to human orthologs described above, 9,637 genes overlapped with human differentially expressed genes (DEGs) and were included in the analysis. Gene expression changes were significantly correlated between the two datasets (R² = 0.389, *P < 0.001*; Figure 2D), suggesting that NT-501 induces broadly similar transcriptional responses *in vivo* and *in vitro*. Notably, 219 genes were commonly differentially expressed in both datasets (adjusted *P < 0.05*). Among these, genes associated with JAK/STAT signaling (red dots in Figure 2B, Figure 2D, and Supplemental Figure 2C) were upregulated, whereas phototransduction-related genes (blue dots in Supplemental Figure 2C) were downregulated.

### 3. CNTF mediates JAK/STAT3 signaling and broad NT-501–induced retinal transcriptional responses

We have demonstrated that NT-501 secretes a broad panel of cytokines and induces a distinct profile of transcriptional responses in retinal tissue (Figure 2, A-D). To determine which retinal tissue responses are specifically mediated by CNTF, we next used a CNTF-specific antibody that selectively neutralizes CNTF-receptor binding (Figure 3A). Western blotting (WB) analysis of hROs treated with Ctrl, NT-501–CM (200 pg/μL CNTF-equivalent), NT-501–CM plus isotype control IgG, or NT-501–CM plus CNTF-neutralizing antibody (nAb) showed that nAb robustly prevented NT-501–CM–induced pSTAT3 accumulation (Figure 3A). To further verify CNTF neutralization, immunohistochemistry (IHC) showed that pSTAT3-positive cells were observed mainly in SOX2-positive Müller glial cells (MGCs), with a subset of PRCs also showing positivity, and this signal was largely abolished by the addition of nAb (Supplemental Figure 3).

**Figure 3.**
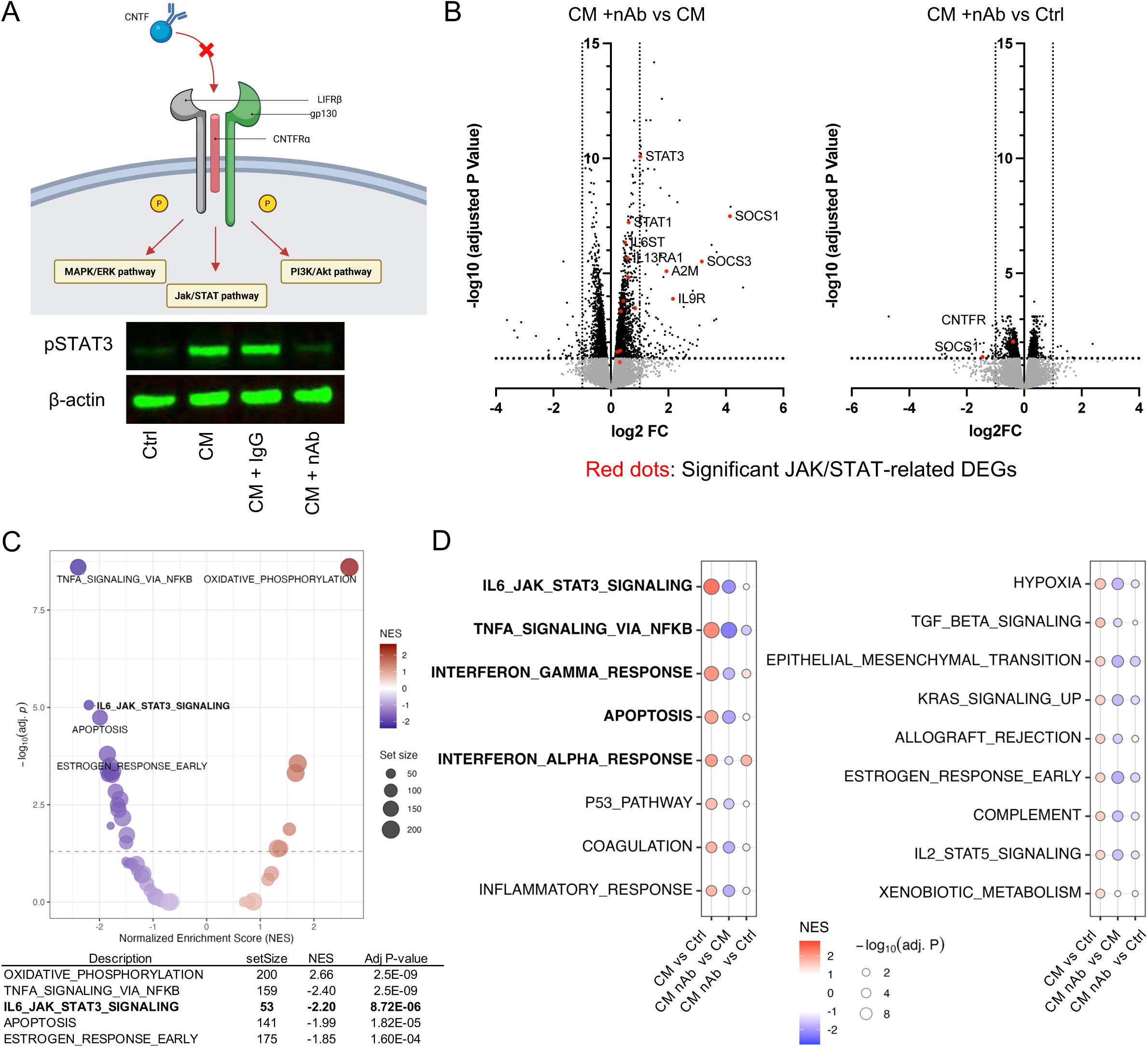
CNTF in NT-501–conditioned medium (NT-501–CM) has the largest effect on changes in gene expression associated with JAK/STAT pathway in hROs. (A) Schematic overview of NT-501–CM and CNTF-neutralizing antibody (nAb) treatment in hROs. Created with BioRender.com. (B) Volcano plots of differentially expressed genes (DEGs) in hROs after 24 hours of treatment, comparing CM plus nAb vs CM (left) and CM plus nAb vs Ctrl medium (right). Black dots indicate DEGs (FDR < 0.05), and red dots highlight JAK/STAT-related genes identified in enrichment analysis. (C) Gene set enrichment analysis (GSEA) of Hallmark gene sets after 24 hours of treatment, showing enrichment in CM + nAb. Each point represents a gene set, plotted by normalized enrichment score (NES) and –log10(adjusted P value). Top 5 significantly enriched gene sets ranked by adjusted P value are shown. (D) Hallmark GSEA dot plot showing selected pathways enriched in NT-501–CM–treated hROs relative to Ctrl after 24 hours of treatment. Dot color indicates normalized enrichment score (NES), and dot size indicates −log10(adjusted P value).

To identify CNTF-dependent transcriptional responses, bulk RNA-seq was performed on NT-501–CM + nAb–treated hROs from the same differentiation batch as the Ctrl-and CM-treated hROs described above (Figure 2), and all three datasets were analyzed together (8 hROs per condition, 3 biological replicates). Differential expression analysis identified 568 upregulated and 539 downregulated genes when comparing CM + nAb with Ctrl, whereas 1,786 upregulated and 1,828 downregulated genes were identified when comparing CM + nAb with CM (adjusted *P < 0.05*; Figure 3B). In volcano plots, JAK/STAT-related gene expression was reduced in CM + nAb relative to CM (Figure 3B), consistent with the WB and IHC findings.

Consistent with this gene-level pattern, Hallmark GSEA of these comparisons showed that IL-6–JAK/STAT3 signaling, TNF-α signaling via NF-κB, and apoptosis were negatively enriched in CM + nAb relative to CM, indicating that these responses are CNTF-dependent (Figure 3C, left). In contrast, the IFN-α and IFN-γ responses remained enriched in CM + nAb relative to Ctrl medium (Figure 3D and Supplemental Figure 4A), suggesting that these responses are largely CNTF-independent. These findings indicate that CNTF mediates a major component of the NT-501–induced retinal transcriptional response, while some residual responses may reflect CNTF-independent cytokine activity from NT-501.

### 4. CNTF attenuates PRC death in a MacTel-relevant degeneration model

We have shown that CNTF mediates a major component of NT-501–induced retinal transcriptional responses, with JAK/STAT signaling representing a prominent CNTF-dependent pathway. CNTF has also been reported to activate additional downstream pathways, including MAPK/ERK and PI3K/AKT signaling (8, 11, 12, 27). WB analysis showed that 4 days of NT-501–CM treatment induced apparent pSTAT3 accumulation in hROs at CNTF-equivalent concentrations of 200 and 1,000 pg/mL, whereas pERK1/2 and pAKT levels showed no apparent changes under the same conditions (Figure 4A). To assess the relative involvement of these downstream signaling pathways in NT-501–CM–treated hROs, we used the JAK1/2 inhibitor baricitinib (Bari) to block JAK-mediated phosphorylation. In NT-501–CM–treated hROs, Bari abolished pSTAT3 induction, whereas pERK1/2 and pAKT levels showed no substantial differences across conditions (Figure 4, B-C).

**Figure 4.**
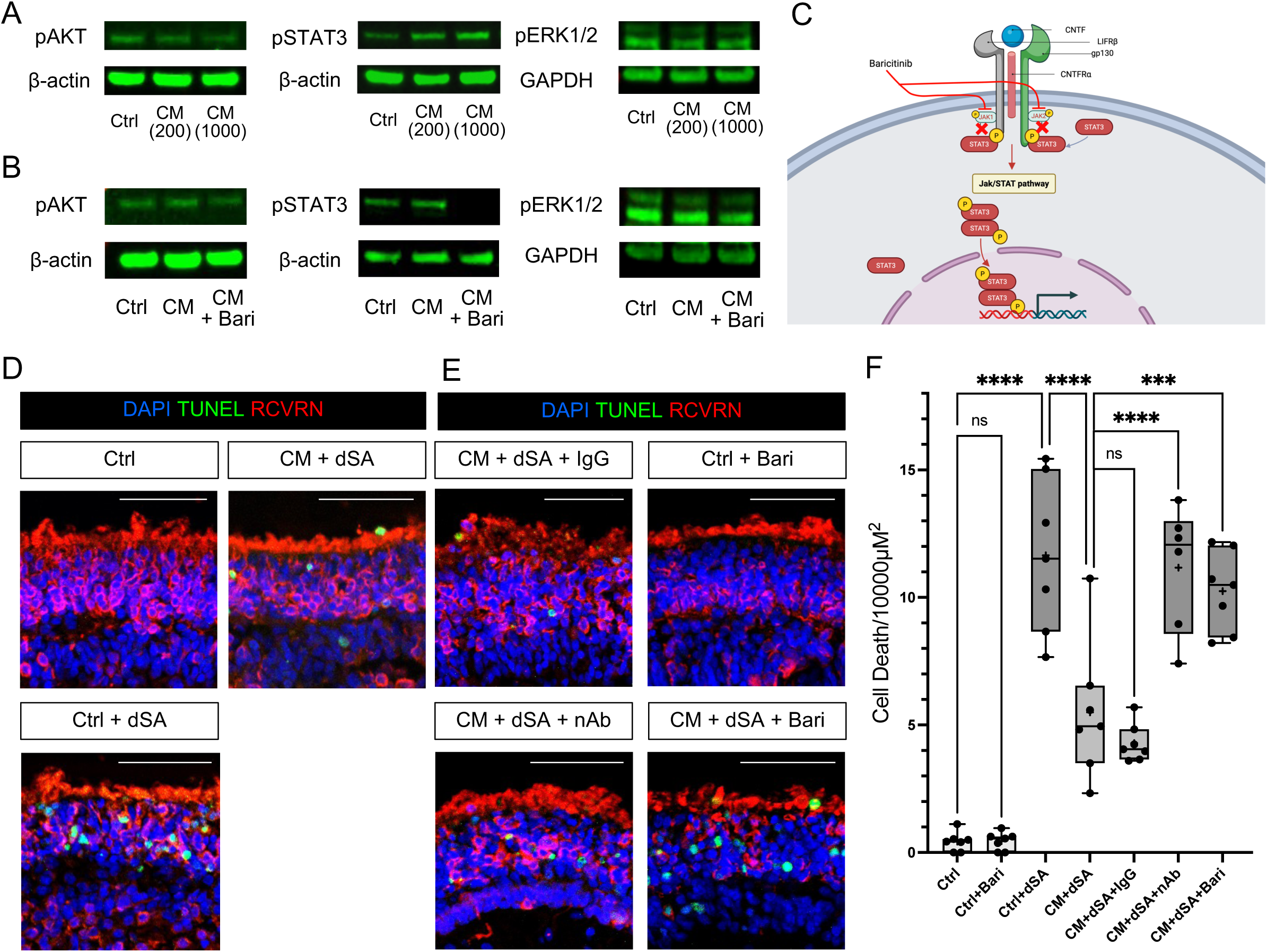
CNTF in NT-501–conditioned medium (NT-501–CM) rescues photoreceptor cell death in 1-deoxysphinganine-toxicity assay. (A-B) Western blot analysis of pERK1/2, pAKT, β-actin, and GAPDH in hROs. β-actin and GAPDH were used as loading controls. Treatment conditions are indicated below the lanes: Ctrl, NT-501–CM at 200 or 1,000 pg/mL CNTF-equivalent in A, and Ctrl, NT-501–CM at 400 pg/mL CNTF-equivalent, or NT-501–CM plus baricitinib (Bari, 2 μM) in B. (C) Schematic of CNTF-induced JAK/STAT signaling and its inhibition by the JAK1/2 inhibitor Bari in hROs. Created with BioRender.com. (D–E) Representative immunostaining images of RCVRN (red; photoreceptors), TUNEL (green; cell death), and DAPI (blue; nuclei) in hROs treated for 4 days. NT-501–CM was adjusted to 1,000 pg/mL CNTF-equivalent. Scale bar, 50 μm. (D) Ctrl (top left), Ctrl + 1-deoxysphinganine (dSA; 1 μM; bottom left), and CM + dSA (top right). (E) CM + dSA + IgG (top left), CM + dSA + nAb (bottom left), Ctrl + Bari (top right), and CM + dSA + Bari (bottom right). (F) Quantification of TUNEL staining from images shown in D and E is presented as box plots. Each dot represents a biologically independent hRO analyzed within the same experiment. Crosses indicate the mean, and horizontal lines indicate the median. Statistical significance was determined using one-way ANOVA followed by Tukey’s multiple-comparisons test. ns, not significant; \*\*\**P < 0.001*; \*\*\*\**P < 0.0001*.

Next, to confirm the therapeutic efficacy of NT-501 in human retinal tissue, we utilized a MacTel-relevant PRC degeneration model in hROs based on 1-deoxysphinganine (dSA)-induced toxicity (26, 28). Prior to the rescue experiments, we confirmed that JAK/STAT-related DEGs were also induced after 4 days of NT-501–CM treatment, and that NT-501–CM-induced gene expression changes were highly correlated between the 1-day and 4-day treatment conditions (10 hROs per condition, 4 biological replicates, Supplemental Figure 4, B-C). We then evaluated whether PRC death could be rescued in the dSA-induced PRC degeneration model. Apoptosis was quantified by TUNEL assay and IHC in hROs treated for 4 days. As previously reported (28), dSA treatment caused significant cell death in PRCs of hROs compared with control treatment; however, treatment with NT-501–CM markedly attenuated PRC death, indicating a functional rescue of dSA-mediated toxicity (Figure 4D).

To test whether CNTF is the therapeutic agent in NT-501–CM, we next neutralized CNTF within NT-501–CM using the CNTF-nAb. We found that the protective effect of NT-501–CM in dSA toxicity was significantly attenuated by CNTF-nAb, whereas this attenuation was not observed with non-specific IgG Ab control (Figure 4E, left, and Figure 4F). Based on pathway analysis in Figure 3, we have shown that CNTF has a broad effect on retinal tissue. To functionally validate the involvement of JAK/STAT signaling specifically, we targeted JAK/STAT signaling using the JAK1/2 inhibitor Bari (Figure 4C). We found Bari was not toxic by itself but significantly attenuated NT-501–CM–mediated rescue of dSA toxicity (Figure 4E, right, and Figure 4F). These data show that NT-501–CM is capable of rescuing cell death in a severe MacTel-relevant toxicity assay, and that this protective effect is largely mediated through CNTF-dependent activation of JAK/STAT signaling.

### 5. Clinically relevant levels of CNTF attenuate photoreceptor cell death in a MacTel-relevant degeneration model

Now that we have identified the CNTF activation of JAK/STAT as the primary therapeutic mechanism in retinal tissue, we next quantitatively assessed the relationship between CNTF levels and both pSTAT3 activation and the PRC rescue effect, using dose–response analysis in hROs. Using a Meso Scale Discovery (MSD) assay, pSTAT3 activation was quantitatively evaluated across a range of CNTF concentrations (25, 50, 100, 200, 400 pg/mL), which included the mean clinically measured vitreous concentration of approximately 50 pg/mL (19). This analysis revealed a linear dose-dependent increase in pSTAT3 in response to CNTF from 0 to 100 pg/ml, reaching a plateau between approximately 100 and 200 pg/mL, with no further increase at higher concentrations (Figure 5A).

**Figure 5.**
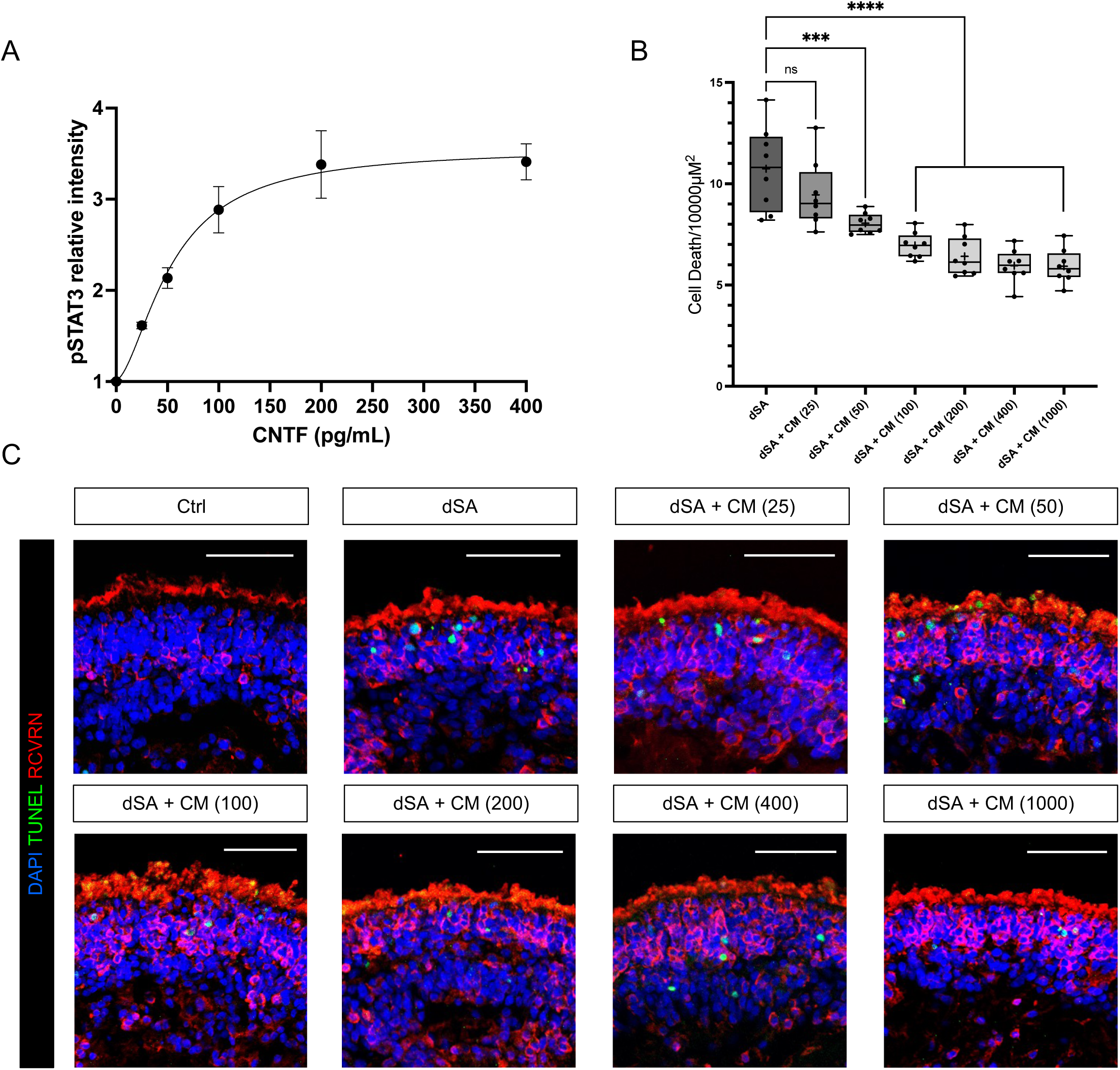
Quantitative analysis of pSTAT3 activity and PRC rescue effects in hROs. (A) Meso Scale Discovery assay of pSTAT3 activity in hROs treated with NT-501–conditioned medium (NT-501–CM) at indicated CNTF-equivalent concentrations. Data are shown as mean ± SEM from two biologically independent experiments. Dose–response curves were fitted using a four-parameter logistic (4PL) model. (B) Quantification of TUNEL staining from images shown in C is presented as box plots. Each dot represents a biologically independent hRO analyzed within the same experiment. Crosses indicate the mean, and horizontal lines indicate the median. Statistical significance was determined using one-way ANOVA followed by Dunnett’s multiple-comparisons test. ns, not significant; \*\*\**P < 0.001*; \*\*\*\**P < 0.0001*. (C) Representative immunostaining images of RCVRN (red; photoreceptors), TUNEL (green; cell death), and DAPI (blue; nuclei) in hROs treated with with Ctrl, dSA, or dSA with NT-501–CM for 4 days. NT-501–CM was adjusted to CNTF-equivalent concentrations of 25, 50, 100, 200, 400, and 1,000 pg/mL. Scale bar, 50 μm.

We next examined whether the dose-dependent changes in STAT3 activation translate into therapeutic efficacy in this model. NT-501–CM was adjusted to CNTF-equivalent concentrations of 25, 50, 100, 200, 400, and 1,000 pg/mL and applied to dSA-treated hROs (8 hROs per condition). Apoptosis of PRCs in the outer nuclear layer was quantified by TUNEL assay and IHC after 4 days of treatment with 1 μM dSA. NT-501–CM significantly reduced PRC death at concentrations ≥50 pg/mL (*P < 0.001*), with stronger effects observed at ≥100 pg/mL (*P < 0.0001*), whereas the effect at 25 pg/mL was not significant (Figure 5, B-C). Notably, median PRC death/10,000 μm² exhibited a dose-dependent reduction that paralleled STAT3 activation, becoming significant at 50 pg/mL and reaching maximal rescue at approximately 200 pg/mL CNTF equivalents in NT-501–CM, with no further improvement at higher concentrations (Figure 5B).

## Discussion

In the present study, we demonstrate three key findings: (i) CNTF acts as the principal mediator of NT-501–induced cytokine signaling, activating the JAK/STAT pathway; (ii) CNTF exhibits dose-dependent effects on STAT3 activation and PRC survival with maximal effects observed at 100-200 pg/mL CNTF; and (iii) our study provides a generalizable framework for defining the mechanisms and optimal dosing of factors released from ECT-based delivery systems.

First, our data show that JAK/STAT signaling is robustly activated across species (rabbit and human), strongly suggesting that NT-501–induced CNTF drives this activation. JAK/STAT activation was observed mainly in MGCs and in a subset of PRCs (11, 29, 30) under CNTF exposure conditions substantially lower than those used in previous supraphysiological models, including genetic overexpression systems (12, 29, 31) and high-dose recombinant CNTF administration (14, 31). In contrast, previous studies also report activation of additional CNTF downstream pathways (e.g., MAPK/ERK and PI3K/Akt) (11, 12, 27, 31), but comparable activation of these pathways was not observed under our experimental conditions. These findings suggest that selective activation of JAK/STAT signaling represents a key mechanism underlying CNTF-mediated neuroprotection at physiologically relevant concentrations.

Second, our experiments demonstrated that both pSTAT3 activation and PRC rescue reached a plateau at approximately 100–200 pg/mL CNTF, with dose-dependent trends observed below this range, suggesting an effective concentration range for CNTF-mediated signaling and PRC rescue (Figure 5, A-C). Notably, the reported vitreous CNTF concentration in patients implanted with NT-501, approximately 50 pg/mL (19), falls within the dose-responsive range observed in our study, suggesting that similar PRC-protective effects may be achieved *in vivo* and may contribute to the slowing of disease progression observed in MacTel patients following NT-501 implantation (22). In addition, metabolic profiling indicated no significant changes in serine-related metabolomics in the rabbit retina with NT-501 (Figure 1C), suggesting that NT-501–mediated PRC protection may not be specific to MacTel and should be broadly applicable to other neurodegenerative diseases. These findings clarify the mechanism of CNTF-mediated PRC protection and identify the effective concentration range, supporting the potential application of this device-based strategy to retinal diseases beyond MacTel.

Finally, this study establishes a generalizable framework that connects experimental models with the physiological conditions associated with NT-501–mediated CNTF delivery in patients. By integrating rabbit *in vivo* implantation with *in vitro* hROs exposed to NT-501–CM, we demonstrate conserved signaling responses across species and systems, including consistent gene expression changes and JAK/STAT pathway activation. Importantly, the use of device-conditioned medium provides a physiologically relevant context that captures the signaling environment generated by the implant, rather than exposure to recombinant protein alone. This experimental configuration enables a systematic approach to defining both mechanism and effective concentration ranges for factors delivered via ECT-based delivery systems in future applications. These findings are therefore likely to reflect retinal responses under clinically relevant conditions and may inform the development of ECT-based strategies for other retinal diseases and therapeutic factors beyond CNTF.

### Limitation of the study

One limitation of this study is that, although we defined the downstream signaling mechanisms induced by clinically relevant CNTF exposure in human retinal tissue and identified CNTF concentrations associated with optimal photoreceptor rescue, the study does not establish how CNTF exposure should be monitored or individualized in patients receiving ECT-based therapy *in vivo*. Because CNTF delivery via ECT devices may vary between individuals, dynamic assessment of CNTF levels in ocular fluids, including aqueous humor and vitreous humor, may become important for optimizing therapeutic efficacy. In this context, approaches enabling real-time estimation or longitudinal monitoring of intraocular CNTF exposure could facilitate patient-specific therapeutic modulation. Together with the mechanistic insights and dose-response relationships presented here, such future efforts may contribute to the establishment of a translational platform for the clinical application of ECT-based therapy.

We demonstrate that JAK/STAT activation through CNTF is the principal mechanism that drives PRC protection in NT-501. This mechanism is conserved across species, as demonstrated in both rabbit retina *in vivo* and hROs *in vitro*, with STAT3 activation in hROs observed predominantly in SOX2-positive MGCs and in a subset of PRCs, supporting both direct and indirect protective effects on PRCs. Quantitative analyses further revealed a concordant dose–response relationship between CNTF exposure, STAT3 activation, and PRC survival, with maximal effects observed within a CNTF-equivalent range of approximately 100–200 pg/mL. These findings define the mechanistic basis of NT-501–mediated PRC protection and identify an optimal concentration range for therapeutic efficacy, providing a framework for translational application and dose optimization in CNTF-based therapies while potentially informing the development of CNTF-based or JAK/STAT–targeted therapeutic strategies for other retinal degenerative diseases.

## Methods

### Sex as a biological variable

Our animal experiments exclusively used male rabbits for RNA-seq and metabolomic analysis. It is unknown whether the findings are relevant for female rabbits.

### Animals

Five adult New Zealand White rabbits (2.59 ± 0.17 kg, range 2.37–2.79 kg) were used. On January 30 in 2024, the NT-501 ECT device (Neurotech Pharmaceuticals) was implanted into the vitreous cavity via pars plana under general anesthesia (in the right eye of each animal). Anesthesia was induced by intramuscular administration of glycopyrrolate (0.1 mg/kg), followed by ketamine (50 mg/kg) after approximately 5 minutes, and xylazine (5 mg/kg) administered approximately 5 minutes later. Animals were monitored for adequate sedation prior to surgery, and the left eyes served as untreated controls. The devices were inserted through the pars plana using a standard surgical approach and positioned within the vitreous cavity. On August 7 in 2024, 190 days after implantation, retinal tissues were collected for downstream analyses. Due to incomplete retinal separation, metabolomic analysis was performed on four rabbits.

### Mass spectrometry analysis

#### Sample preparation and metabolite extraction

Metabolites were extracted from retinal tissue samples using a modified biphasic methanol/water/chloroform extraction procedure based on a previously described method (32). Approximately 20 mg of tissue was transferred to a 2 mL Eppendorf tube and spiked with a stable isotope-labeled amino acid internal standard mixture (Supplemental Table 2). Methanol and water were subsequently added, and samples were homogenized using 1.4 mm ceramic beads in a Retsch MM400 homogenizer for 2 min at 25 Hz. Following homogenization, the supernatant was transferred to a fresh 2 mL Eppendorf tube to remove the ceramic beads, and chloroform was subsequently added to generate a biphasic extraction system. Phase separation was achieved by centrifugation, after which the lower organic phase was collected for further processing. The remaining aqueous/methanol layer was subjected to a second extraction using formic acid and chloroform. The aqueous/methanol extract was retained for amino acid analysis and stored at -80°C until LC–MS/MS analysis.

### LC–MS/MS Analysis

LC–MS/MS analyses was performed using a Waters Acquity UPLC system coupled to a Waters TQ Absolute triple quadrupole mass spectrometer operated in positive electrospray ionization mode. Mass spectrometric detection was performed in scheduled multiple reaction monitoring mode using MassLynx version 4.2 software (Waters Corporation). Relative quantification was performed using peak area ratios of analytes to their corresponding stable isotope-labeled internal standards. Data processing and peak integration were performed using TargetLynx XS software.

Amino acid separation was achieved using an Astec Chirobiotic T column (5 µm, 150 × 4.6 mm) maintained at 25°C. The autosampler temperature was set to 10°C and the injection volume was 1 µL. Mobile phase A consisted of water containing 0.1% formic acid, while mobile phase B consisted of methanol. Chromatographic separation was performed using isocratic elution at a flow rate of 0.35 mL/min over a total run time of 25 min. Source parameters were set as follows: capillary voltage, 0.7 kV; source temperature, 150°C; desolvation temperature, 500°C; cone gas flow, 150 L/h; and desolvation gas flow, 900 L/h.

### Human iPSC maintenance

The human SMT4C1 iPSC line was derived by the Salk Institute iPSC Core Facility using peripheral blood mononuclear cells from a male donor. Reprogramming was performed using Sendai virus–based system for delivery of reprogramming factors. The cell line was confirmed to have a normal karyotype and was routinely tested for mycoplasma contamination. iPSCs were maintained on Cultrex (R&D Systems, 3432-005-01)-coated plates with mTeSR Plus medium (STEMCELL Technologies, 100-0276). Cells were passaged every 3–4 days at approximately 80% confluence using EDTA-based dissociation. Colonies containing clearly visible differentiated cells were marked and manually removed before passaging.

### Differentiation of hROs

To make hROs, embryoid bodies were formed (day 0) using Aggrewells (STEMCELL Technologies, 34811) in Neural Induction Medium containing DMEM/F12 (Gibco, 11330057) with 1% N2 supplement (Gibco, 17502048), 1% MEM NEAA (Gibco, 11140050), 2 mg/mL heparin (STEMCELL Technologies, 07980) at 180 U/mg, and 1% Pen-Strep (Gibco, 15140122) and then plated (day 7) onto Matrigel growth factor reduced (Corning, 47743-718) coated plates. Media was switched to Retinal Differentiation Medium (RDM) containing 48% DMEM/F12 and 48% DMEM (Gibco, 11995073) supplemented with 2% B27 supplement without vitamin A (Gibco, 12587010), and 1% MEM NEAA on day 16. On day 28, plated EBs were fragmented using a sterile scalpel (McKesson, 801453), scraped with a P1000 pipette, collected, and transferred to suspension culture on an orbital shaker (130 rpm). At week 8, mature organoids were transferred to a bioreactor (35 rpm), and the medium was changed to RDM with 10% FBS (Corning, 35-016-CV), 100 μM Taurine (Sigma, TO-625), and 2 mM Glutamax (Gibco, 35050061)10% FBS, 100 μM taurine, and 2 mM Glutamax. The remainder of the differentiation protocol is as previously published (33, 34). hROs were transferred from the bioreactor at week 17 and plated in low attachment 6 well plates.

### NT-501–conditioned medium

Implants were maintained in primary jars containing human endothelial serum-free medium (Ctrl; Neurotech Pharmaceuticals) at 37°C for 4 weeks. The implants were then removed from the primary jars, and the medium from each jar was collected separately without pooling (CM-1 to CM-4). The implants were washed once with Ca^2+^/Mg^2+^-free HBSS and individually incubated in 1 mL Ctrl medium in separate wells of a 24-well tissue culture plate for 24 hours. After incubation, the media from each well were pooled and stored at −80°C (CM-5). Both primary jar-derived conditioned media (CM-1 to CM-4) and pooled 24-hour incubation conditioned medium (CM-5) were defined as NT-501–CM, and CNTF concentrations were quantified using the Quantikine CNTF ELISA kit (R&D Systems, DNT00).

### Multiple cytokine assay

Cytokine concentrations in Ctrl and NT-501–CM were measured using the Human Cytokine/Chemokine 96-Plex Discovery Assay (Eve Technologies Corporation, HD96), which uses Luminex xMAP technology to simultaneously quantify 96 cytokines, chemokines, and growth factors. A total of six independent samples (Ctrl and CM-1 through CM-5) were assayed in triplicate without dilution (dilution factor = 1). The complete list of analytes and their assay sensitivities is provided in Supplemental Table 1.

### RNA-seq analysis

Total RNA was extracted from rabbit retinas and hROs using TRIzol Reagent (Thermo Fisher, 15596026) and purified using the RNeasy Plus Micro Kit (Qiagen, 74034) according to the manufacturers’ instructions. For each sample, 100 ng of total RNA was used to prepare sequencing libraries using the NEBNext Ultra II RNA Library Prep Kit with poly(A) selection (New England Biolabs) according to the manufacturer’s recommended protocol. Libraries were dual-indexed using NEBNext Multiplex Oligos for Illumina (Dual Index Primers Set 2; New England Biolabs, E7780S) and sequenced on an Element AVITI platform using paired-end sequencing.

Quality control of FastQ files was performed using FastQC (v0.12.0) and MultiQC (35). Reads were mapped to the Ensembl reference *Oryctolagus cuniculus* genome using Rsubread (v2.12) function *align*. Mapped reads were then counted using Rsubread *featureCounts* function. Lowly expressed genes were filtered using the limma (v3.58) function *filterByExpr*. Non-protein-coding genes were also discarded. Data was normalized using TMM normalization (36). Sample quality was assessed by multidimensional scaling analysis. Differential expression was performed using the limma-voom pipeline (37). Quality weights for each sample were estimated using the limma function *voomWithQualityWeights*. Correction for the false discovery rate was performed using the Benjamini-Hochberg correction, and an adjusted P value less than 0.05 was considered significant for differential expression.

Rabbit DEGs were mapped to human orthologs using Ensembl BioMart via the biomaRt package (version 2.66.1). Queries were performed against the *Oryctolagus cuniculus* dataset, retaining only high-confidence human ortholog mappings. Cross-species comparisons between rabbit and human datasets were performed using human ortholog–mapped genes, and correlations were assessed by Pearson correlation analysis and simple linear regression. GSEA was performed using the clusterProfiler package (version 4.18.4), with genes ranked by log2FC. KEGG gene sets were obtained through the clusterProfiler, and Hallmark gene sets were obtained from MSigDB via the msigdbr package (version 26.1.0). Rabbit GSEA was performed with the human ortholog-mapped genes for compatibility with these human gene sets. Volcano and enrichment plots were generated using ggplot2 (version 4.0.2) and enrichplot (version 1.30.5), respectively. All analyses were conducted in R (version 4.5.2).

### Western Blotting

hRO lysates (6-10 hROs per condition) were prepared using RIPA buffer (ThermoFisher, 89901) with an added protease and phosphatase inhibitor (ThermoFisher, A32961). Lysates were incubated on ice for 30 minutes and sonicated for 10 seconds × 3 cycles, incubating on ice for 50 seconds after each sonication. The lysates were then centrifuged at 20,000 × g for 10 minutes at 4°C and the supernatant was stored at -80°C. The total protein concentration was calculated using a bicinchoninic acid protein assay (Thermo Fisher Scientific, 23227). Lysates were separated by SDS-PAGE gel (Thermo Fisher Scientific, nw04125box) and transferred to a nitrocellulose membrane. The membrane was blocked for 1 hour and incubated overnight at 4°C with primary antibodies diluted as listed in Supplemental Table 3 in 5% milk/TBST. After washing with TBST, the membrane was then incubated with a fluorescent secondary antibody (1:20,000, LICOR, 926-32212, 926-32213) in 5% milk buffer for 2 hours at room temperature (RT) and imaged using the LI-COR Odyssey imaging system.

### Meso Scale Discovery (MSD) assay

All samples were analyzed using the MSD Phospho-STAT3 (Tyr705) Kit (K150SVD-1). Protein samples were collected by removing hROs from conditioned media and rinsing twice with PBS. Non-retinal regions were removed prior to tissue collection. Samples were flash-frozen in liquid nitrogen and stored at −80°C. hRO lysates were prepared using MSD lysis buffer. Lysates were incubated on ice for 30 minutes and sonicated (10 seconds × 3 cycles) with 50-second intervals on ice between cycles. Lysates were centrifuged at 20,000 × g for 10 minutes at 4°C, and the supernatant was collected and stored at −80°C. A total of 20 μg protein per well was loaded onto a 96-well MSD plate pre-coated with capture antibodies. Plates were incubated at RT with shaking at 500 rpm for 1 hour, followed by three washes with 1× wash buffer. Detection antibodies (25 μL per well) were added and incubated at RT with shaking at 500 rpm for 1 hour. Read buffer was added immediately prior to signal detection using the MESO QuickPlex SQ 120. Data were pooled from two biologically independent experiments (n = 10 and 36 hROs per condition, respectively).

### dSA-induced PRC degeneration assay in hROs

To assess PRC death, we employed a MacTel-specific PRC degeneration model in hROs based on dSA-induced toxicity, using 1-deoxysphinganine (dSA; Avanti Research, 860493). Seven to eight hROs were assigned to each condition: control, dSA-treated, and dSA-treated conditions with NT-501–CM, in the presence or absence of isotype control IgG (R&D Systems, AB-108-C), nAb, (R&D Systems, AF-257-NA) or the JAK1/2 inhibitor Bari (APExBIO, A4141). In dose–response experiments, NT-501–CM was applied at varying CNTF-equivalent concentrations as indicated. nAb and Bari were dissolved in PBS (Thermo Fisher, 14190250) and dimethyl sulfoxide (DMSO) to generate 200 μg/mL and 10 mM stock solutions, respectively. NT-501–CM and antibodies (IgG or nAb) were preincubated overnight at 37℃ prior to application. hROs were pretreated with an inhibitor-containing medium for 1 hour, followed by treatment with dSA and NT-501–CM under the indicated conditions. Treatments were maintained for 4 days, with full medium replacement including dSA every 48 hours. In contrast, NT-501–CM, antibodies, and inhibitors were added daily. Final treatment conditions included dSA (1 μM), NT-501–CM (adjusted to the indicated CNTF-equivalent concentrations), antibodies (IgG or nAb; molar ratio 1:1500 relative to CNTF), and Bari (2 μM), as appropriate. Vehicle controls (ethanol, PBS, and DMSO) were included where applicable. After 4 days of treatment, organoids were collected for downstream analyses. PRC death was assessed by TUNEL assay and IHC, and protein expression was analyzed by WB analysis.

### Immunohistochemistry and TUNEL assay

The hROs were fixed with 4% paraformaldehyde (Thermo Scientific, 28906) for 15 minutes at RT, infiltrated with 15% sucrose/PBS for 15 minutes at RT, 30% sucrose/PBS overnight at 4°C, embedded in O.C.T. compound (Fisher Healthcare, 4585), and stored at -80°C until sectioning. Cryosections were cut at 14-μm thickness using a cryostat OTF7000 (Bright Instruments). Sections were washed with PBS and blocked with a blocking solution of 5% donkey serum containing 0.1% Triton X-100 in PBS for 1 hour at RT. Primary antibodies were diluted to the dilution listed in Supplemental Table 3 and incubated overnight at 4°C. Sections were then washed three times with PBS, and incubated with the appropriate secondary antibodies and DAPI (Invitrogen, D1306) for 2 hours at RT protected from light. After secondary antibody staining, the samples were washed three times with PBS and mounted using Prolong Glass Antifade Mountant (Invitrogen, 3224967). With regard to pSTAT3 staining, we added phosphatase inhibitor (Thermo Scientific, A32957) to the fixation and subsequent blocking steps. Antigen unmasking was performed by submerging tissue sections in 1X Signal Stain EDTA Unmasking Solution (Cell Signaling Technology, 14747) diluted in water and heated to 95°C for 15 minutes before blocking. Stained samples were kept in the dark and stored at 4°C until imaging with a confocal microscope (Zeiss, LSM710).

TUNEL staining was performed using In Situ Cell Death Detection Kit, Fluorescein (Sigma-Aldrich, 11684795910) according to the manufacturer’s instructions before primary antibody incubation. Each hRO was represented by one central cryostat slice. Overlapping TUNEL-positive and DAPI-positive signals within RCVRN-positive cells in the outer nuclear layer were counted as photoreceptor cell death. Cell death was normalized to the area of RCVRN-staining in each hRO, as previously reported (34). Images used for quantification were acquired using identical laser power, gain, offset, and exposure settings across conditions. For each condition, 7–8 hROs from a single differentiation batch were analyzed. Similar results were obtained in an independent differentiation batch.

### Statistics

Statistical analyses for non–RNA-seq data were performed using Prism 10 (GraphPad). Experiments were performed at least twice independently to confirm reproducibility. Replicate numbers are indicated in the corresponding figure legends. Paired t tests with Holm–Šídák correction were used to compare implanted and contralateral control eyes in rabbits. Differences among three or more groups were assessed using one-way ANOVA with Tukey’s or Dunnett’s multiple-comparison tests, as indicated in the figure legends. *P* < *0.05* was considered statistically significant.

## Study approval

All animal procedures were approved by the Scripps Research Institute’s Institutional Animal Care and Use Committee (IACUC), and were conducted in accordance with federal animal experimental guidelines.

## Data availability

RNA-seq data will be made available upon publication. All analyses were performed using publicly available R packages as described in the Methods.

## Author’s contributions

Designing and conducting experiments: Y.I., L.L., S.G., A.M., S.H.P., K.T., M.L.G., and K.T.E.; Analyzing data: Y.I., L.L., M.V.D., and R.B.; Interpreting data: Y.I., M.L.G., and K.T.E.; Writing—original draft: Y.I., with contributions from L.L., S.G., A.M., K.T., M.V.D., R.B. to the Methods section; Writing—review & editing: S.G., K.N., M.L.G., M.F., and K.T.E., Supervision: M.L.G., M.F., and K.T.E.; Funding acquisition: M.F.

## Acknowledgement

We would like to thank the Lowy family for their longstanding support of the Lowy Medical Research Institute and the MacTel Project. The human iPSC-SMT4C1 cell line was provided by the Salk Institute iPSC Core Facility. We thank Neurotech Pharmaceuticals (Cumberland, RI, USA), particularly Lisa Orecchio, Arne Nystuen, and Konrad Kauper, for providing NT-501–conditioned medium and for experimental discussions. We thank Edith Aguilar and Ray Gariano for supporting the rabbit experiments. We thank J. Shimashita and S. Head at TSRI DNA Array Core for their work on RNA-Seq. Y.I. was supported by grants from the Manpei Suzuki Diabetes Foundation.

## Supplemental Information

**Supplemental Figure 1.**
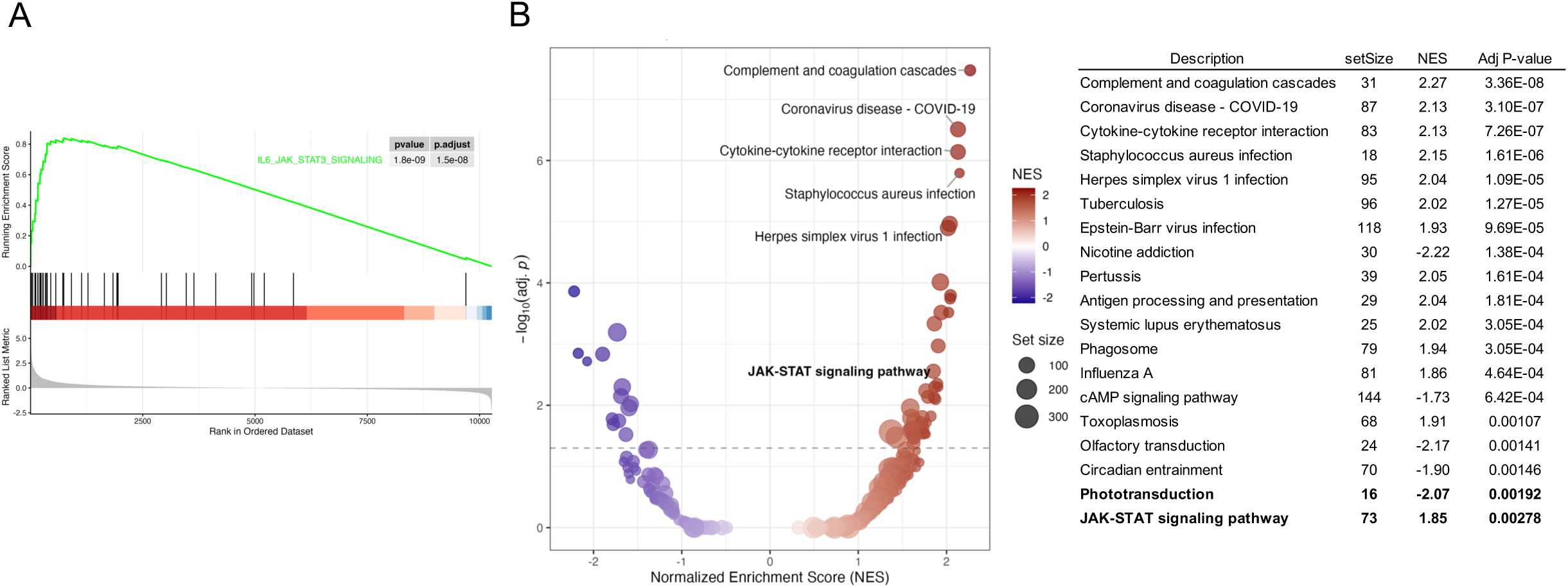
Hallmark and KEGG GSEA results for NT-501–treated rabbit retinas relative to contralateral control eyes (A) Hallmark enrichment plot for the IL-6–JAK/STAT3 signaling gene set in rabbit retinas following NT-501 implantation compared with contralateral controls. (B) KEGG pathway GSEA showing enrichment in NT-501–treated rabbit retinas relative to contralateral control eyes.

**Supplemental Figure 2.**
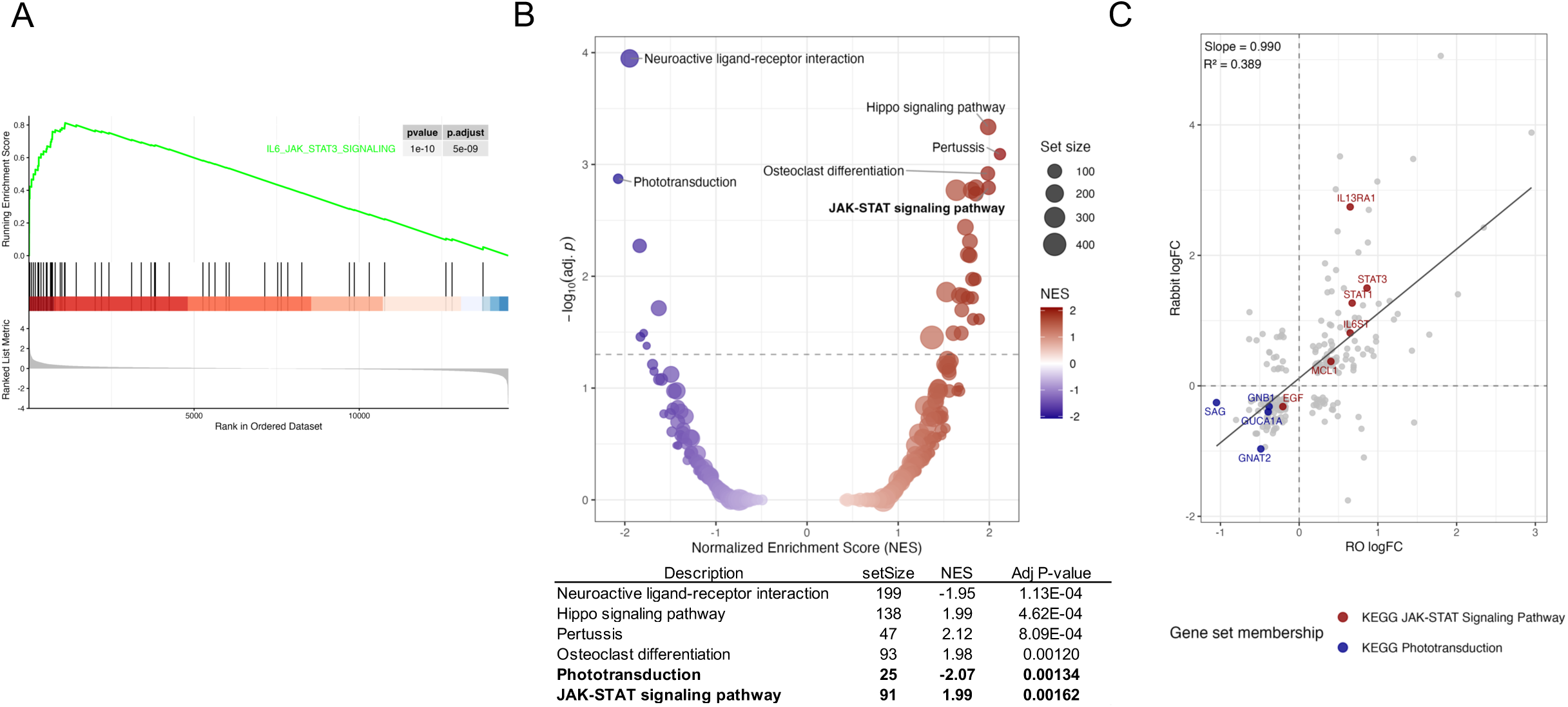
Supplemental transcriptional responses of hROs treated with NT-501–conditioned medium (NT-501–CM) for 24 hours (A) Hallmark enrichment plot for the IL-6–JAK/STAT3 signaling gene set in NT-501–CM relative to Ctrl. (B) KEGG pathway GSEA of NT-501–CM–treated hROs relative to Ctrl. (C) Comparison of gene expression changes between NT-501–implanted rabbit retina and NT-501–CM–treated hROs after 24 hours of treatment. Red and blue dots indicate IL-6–JAK/STAT3 signaling and phototransduction genes, respectively. The solid line indicates linear regression, with slope and R² shown.

**Supplemental Figure 3.**
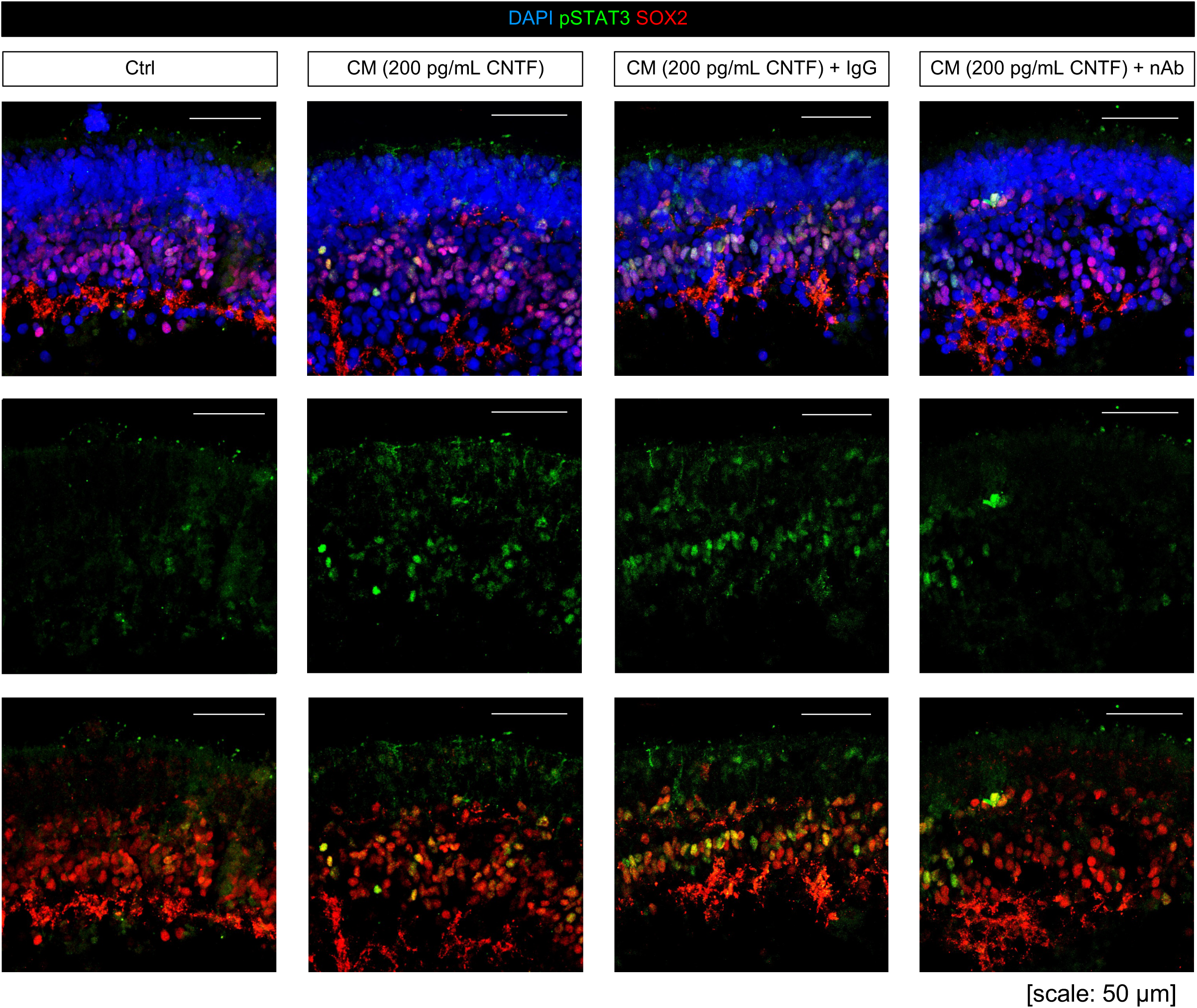
Representative immunostaining images of pSTAT3 (green), SOX2 (red), and DAPI (blue) in hROs treated with control medium, NT-501–conditioned medium (NT-501–CM) adjusted to 200 pg/mL CNTF-equivalent, NT-501–CM plus isotype control IgG, or NT-501–CM plus CNTF-neutralizing Ab.

**Supplemental Figure 4.**
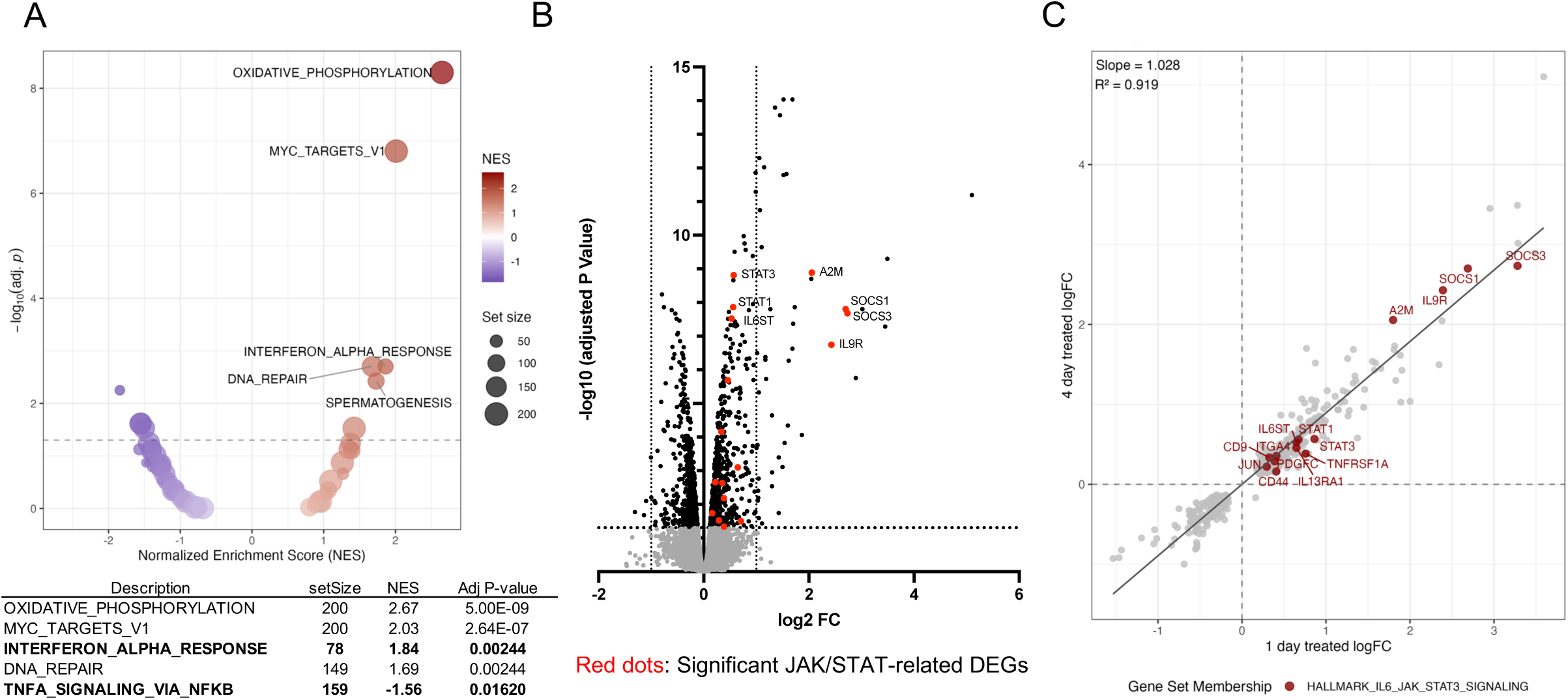
Supplemental transcriptional analyses of NT-501–conditioned medium (NT-501–CM)–treated hROs (A) Hallmark enrichment plots for the IL-6–JAK/STAT3 signaling gene set after 24 hours of treatment: NT-501–CM + nAb relative to Ctrl. (B) Volcano plots of differentially expressed genes (DEGs) in hROs after 4 days of NT-501–CM treatment compared with Ctrl medium. Black dots indicate DEGs (FDR < 0.05), and red dots highlight JAK/STAT-related genes identified in enrichment analysis. (C) Comparison of gene expression changes between 1 day and 4 days of NT-501–CM treatment. Red and blue dots indicate JAK/STAT signaling and phototransduction genes, respectively. The solid line indicates linear regression, with slope and R² shown.

**Supplemental Table 1.** Multiplex cytokine profiling of five independent NT-501–conditioned media samples (Ctrl and CM-1 through CM-5)

**Supplemental Table 2.** Stable Isotope-Labeled Internal Standards Used for Amino Acid Analysis

**Supplemental Table 3.** Antibodies for Western Blotting and Immunohistochemistry

## References

1. Orive G, et al. Encapsulated cell technology: from research to market. Trends Biotechnol. 2002;20(9):382–387.

2. Santos-Vizcaino E, et al. Clinical Applications of Cell Encapsulation Technology. Methods Mol Biol. 2020;2100(473–491.

3. Wang J, et al. Encapsulated cell technology: Delivering cytokines to treat posterior ocular diseases. Pharmacol Res. 2024;203(107159.

4. Curtis R, et al. Retrograde axonal transport of ciliary neurotrophic factor is increased by peripheral nerve injury. Nature. 1993;365(6443):253–255.

5. Wen R, et al. CNTF and retina. Prog Retin Eye Res. 2012;31(2):136–151.

6. Pasquin S, et al. Ciliary neurotrophic factor (CNTF): New facets of an old molecule for treating neurodegenerative and metabolic syndrome pathologies. Cytokine Growth Factor Rev. 2015;26(5):507–515.

7. Shpak AA, et al. Ciliary neurotrophic factor in patients with primary open-angle glaucoma and age-related cataract. Mol Vis. 2017;23(799–809.

8. Nystuen A, et al. Neuroprotective properties of ciliary neurotrophic factor in the retina for the treatment of macular telangiectasia type 2. Cytokine Growth Factor Rev. 2025;84(12–19.

9. Dittrich F, et al. Ciliary neurotrophic factor: pharmacokinetics and acute-phase response in rat. Ann Neurol. 1994;35(2):151–163.

10. Wen R, et al. Injury-induced upregulation of bFGF and CNTF mRNAS in the rat retina. J Neurosci. 1995;15(11):7377–7385.

11. Peterson WM, et al. Ciliary neurotrophic factor and stress stimuli activate the Jak-STAT pathway in retinal neurons and glia. J Neurosci. 2000;20(11):4081–4090.

12. Rhee KD, et al. Molecular and cellular alterations induced by sustained expression of ciliary neurotrophic factor in a mouse model of retinitis pigmentosa. Invest Ophthalmol Vis Sci. 2007;48(3):1389–1400.

13. Pease ME, et al. Effect of CNTF on retinal ganglion cell survival in experimental glaucoma. Invest Ophthalmol Vis Sci. 2009;50(5):2194–2200.

14. Bucher F, et al. CNTF Prevents Development of Outer Retinal Neovascularization Through Upregulation of CxCl10. Invest Ophthalmol Vis Sci. 2020;61(10):20.

15. Do Rhee K, et al. Ciliary neurotrophic factor-mediated neuroprotection involves enhanced glycolysis and anabolism in degenerating mouse retinas. Nat Commun. 2022;13(1):7037.

16. Tao W, et al. Encapsulated cell-based delivery of CNTF reduces photoreceptor degeneration in animal models of retinitis pigmentosa. Invest Ophthalmol Vis Sci. 2002;43(10):3292–3298.

17. Emerich DF, and Thanos CG. NT-501: an ophthalmic implant of polymer-encapsulated ciliary neurotrophic factor-producing cells. Curr Opin Mol Ther. 2008;10(5):506–515.

18. Zhang K, et al. Ciliary neurotrophic factor delivered by encapsulated cell intraocular implants for treatment of geographic atrophy in age-related macular degeneration. Proc Natl Acad Sci U S A. 2011;108(15):6241–6245.

19. Kauper K, et al. Two-year intraocular delivery of ciliary neurotrophic factor by encapsulated cell technology implants in patients with chronic retinal degenerative diseases. Invest Ophthalmol Vis Sci. 2012;53(12):7484–7491.

20. Birch DG, et al. Long-term Follow-up of Patients With Retinitis Pigmentosa Receiving Intraocular Ciliary Neurotrophic Factor Implants. Am J Ophthalmol. 2016;170(10–14.

21. Goldberg JL, et al. Phase I NT-501 Ciliary Neurotrophic Factor Implant Trial for Primary Open-Angle Glaucoma: Safety, Neuroprotection, and Neuroenhancement. Ophthalmol Sci. 2023;3(3):100298.

22. Chew EY, et al. Cell-Based Ciliary Neurotrophic Factor Therapy for Macular Telangiectasia Type 2. NEJM Evid. 2025;4(8):EVIDoa2400481.

23. Charbel Issa P, et al. Macular telangiectasia type 2. Prog Retin Eye Res. 2013;34(49–77.

24. Powner MB, et al. Loss of Müller’s cells and photoreceptors in macular telangiectasia type 2. Ophthalmology. 2013;120(11):2344–2352.

25. Scerri TS, et al. Genome-wide analyses identify common variants associated with macular telangiectasia type 2. Nat Genet. 2017;49(4):559–567.

26. Gantner ML, et al. Serine and Lipid Metabolism in Macular Disease and Peripheral Neuropathy. N Engl J Med. 2019;381(15):1422–1433.

27. Kassen SC, et al. CNTF induces photoreceptor neuroprotection and Müller glial cell proliferation through two different signaling pathways in the adult zebrafish retina. Exp Eye Res. 2009;88(6):1051–1064.

28. Rosarda JD, et al. Imbalanced unfolded protein response signaling contributes to 1-deoxysphingolipid retinal toxicity. Nat Commun. 2023;14(1):4119.

29. Rhee KD, et al. CNTF-mediated protection of photoreceptors requires initial activation of the cytokine receptor gp130 in Müller glial cells. Proc Natl Acad Sci U S A. 2013;110(47):E4520–4529.

30. Lipinski DM, et al. CNTF Gene Therapy Confers Lifelong Neuroprotection in a Mouse Model of Human Retinitis Pigmentosa. Mol Ther. 2015;23(8):1308–1319.

31. Wang Y, et al. Impacts of ciliary neurotrophic factor on the retinal transcriptome in a mouse model of photoreceptor degeneration. Sci Rep. 2020;10(1):6593.

32. Cordes T, and Metallo CM. Quantifying Intermediary Metabolism and Lipogenesis in Cultured Mammalian Cells Using Stable Isotope Tracing and Mass Spectrometry. Methods Mol Biol. 2019;1978:219–241.

33. Cowan CS, et al. Cell Types of the Human Retina and Its Organoids at Single-Cell Resolution. Cell. 2020;182(6):1623–1640.e1634.

34. Eade K, et al. Toxicity Screens in Human Retinal Organoids for Pharmaceutical Discovery. J Vis Exp. 2021;169:e62269.

35. Ewels P, et al. MultiQC: summarize analysis results for multiple tools and samples in a single report. Bioinformatics. 2016;32(19):3047–3048.

36. Robinson MD, and Oshlack A. A scaling normalization method for differential expression analysis of RNA-seq data. Genome Biol. 2010;11(3):R25.

37. Law CW, et al. RNA-seq analysis is easy as 1-2-3 with limma, Glimma and edgeR. F1000Res. 2018;5:1408.

